# Membrane-mediated interactions induce spontaneous filament bundling

**DOI:** 10.1101/336545

**Authors:** Afshin Vahid, George Dadunashvili, Timon Idema

## Abstract

The plasma membrane and cytoskeleton of living cells are closely coupled dynamical systems. Internal cytoskeletal elements such as actin filaments and microtubules continually exert forces on the membrane, resulting in the formation of membrane protrusions. In this paper we investigate the interplay between the shape of a cell distorted by pushing and pulling forces generated by microtubules and the resulting rearrangement of the microtubule network. From analytical calculations, we find that two microtubules that deform the vesicle can both attract or repel each other, depending on their angular separations and the direction of the imposed forces. We also show how the existence of attractive interactions between multiple microtubules can be deduced analytically, and further explore general interactions through Monte Carlo simulations. Our results suggest that the commonly reported parallel structures of microtubules in both biological and artificial systems can be a natural consequence of membrane mediated interactions.

## 1 Introduction

Cells are enveloped by a plasma membrane which serves as a selective soft physical barrier and is home to many functional proteins. The stability and shape of cellular membranes are determined not only by inherent properties of the membrane, but also by interactions with the cell’s cytoskeleton (1). The highly dynamic cytoskeletal network is vital for numerous biological processes, including cell motility, cell migration, and cell signaling (2, 3). A typical feature occurring in such processes is the formation of membrane protrusions. Protrusions commonly emerge in the form of microvilli, filopodia or lamellipodia (4, 5). These leading-edge protrusions, the existence of which is vital for responding to external cues, can be driven, controlled and elongated by a complicated crosstalk between the membrane and underlying filaments.

The spatial arrangement of cytoskeletal filaments, force generation mechanisms, and cytoskeletal networks coupling to the shape of cells have been investigated extensively, both theoretically and experimentally (6–11). For example, when growing encapsulated microtubules inside an artificial spherical membrane, it has been shown that the vesicle exhibits a diverse range of morphologies, from a simple elongated shape to dumbbell-like geometries (7). The diversity in the shape of such vesicles results from both the elongation dynamics of the filaments inside them and the material properties of the membrane. Such spatial rearrangement of filaments stems from the conditions imposed on them from various elements, one of which is the cell shape.

In this paper, we investigate the interplay between the shape of vesicles that are deformed by internal force generating filaments like microtubules, and the rearrangement of those filaments. In a biological cell, microtubules undergo treadmilling and dynamic instabilities (catastrophes) which are controlled by associated proteins (12). Only a few of the microtubules that grow inside a cell can reach the cell membrane (13). The pushing and pulling forces generated by those few microtubules can be harnessed for creating protrusions of the membrane (14). Membrane mediated interactions between microtubule-induced protrusions may influence the arrangement of other functional filaments in addition to microtubules themselves (15, 16). Therefore, we study how the presence of a lipid bilayer membrane, which has both elastic and fluid properties, alters the interaction between microtubules. This interaction could both drive processes like the formation of filament bundles or inhibit microtubule aggregation.

We use a modified version of the theoretical framework that has been developed for investigating membrane mediated interactions between proteins embedded in or bounded to a fluid membrane (17, 18). We first explain the model in detail. We then study the effects of all the possible elements on the interaction between microtubules. We reveal that the relative orientation of the deforming forces (generated by the microtubules) determines the nature of their interactions. In particular, we demonstrate that bundling is a stable effect observed for any number of interacting microtubules with forces of the same orientation. Our results thus elucidate the effective role of the membrane in determining the equilibrium arrangement of protrusions imposed by the cytoskeleton.

## 2 Model

Our model is based on minimizing the energy of the system consisting of the membrane and force imposing microtubules, depicted in figure 1, to obtain the equilibrium shape. We use the Canham-Helfrich functional (19, 20)to characterize the energy, due to elastic deformations of a closed membrane vesicle.

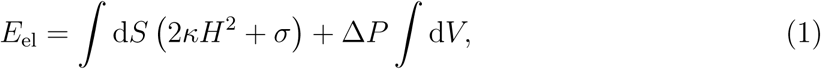

with *H, κ, σ* and Δ*P* the mean curvature, bending modulus, surface tension and the pressure difference (between inside and outside of the vesicle), respectively. The first term in the elastic energy penalizes high curvature, while the second and third terms penalize the change in projected surface area and enclosed volume of the vesicle. In order to describe microtubules deforming the membrane, we impose deformations at *n* points **r**(*θ*_*i*_, *ϕ*_*i*_) as boundary conditions

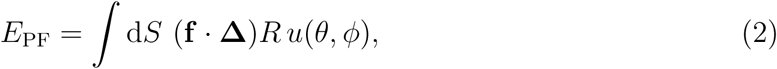

where **f** ∈ ℝ^*n*^ is the vector of forces, exerted on the membrane by the microtubules. This vector contains the projections of the point forces along the radial direction of a spherical vesicle. We denote by **Δ** the vector of Dirac-delta functions at the points (*θ*_*i*_, *ϕ*_*i*_),

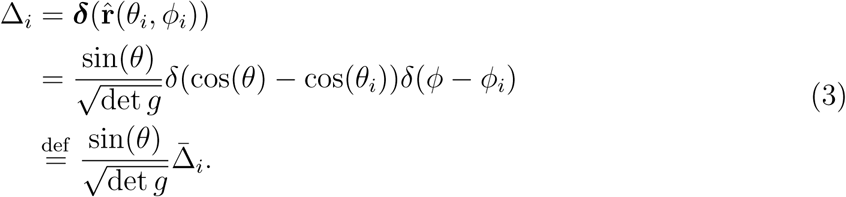

Here *g* is the metric tensor of the membrane, defined as *g*_*μν*_ = *∂*_*μ*_***r*** · *∂*_*ν*_***r***. The metric also appears in the surface area element 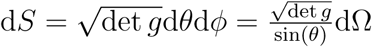.

**Figure 1:**
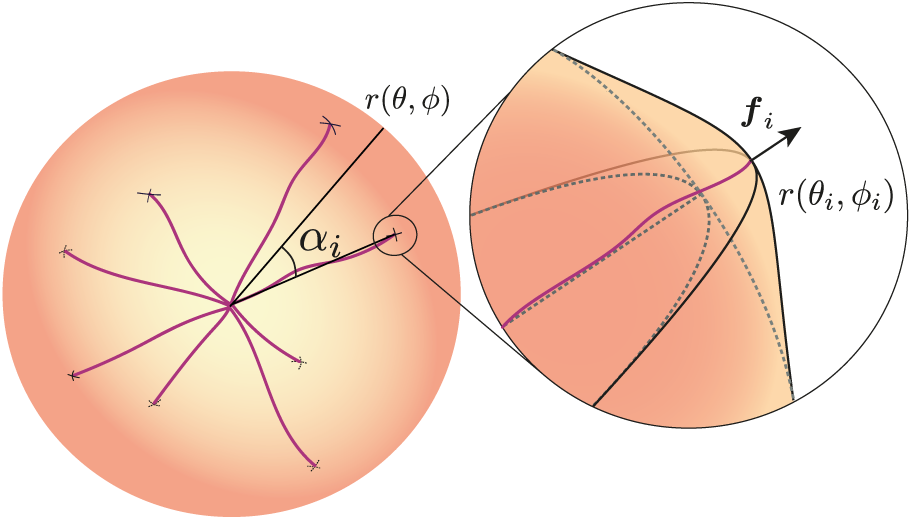
A schematic representation of a spherical membrane vesicle containing microtubules. **Inset**: local (small) deformation *R u*(*θ*_*i*_, *ϕ*_*i*_), caused by a microtubule, which is exerting a force of magnitude *f*_*i*_, along the radius of the vesicle.

**Figure 2:**
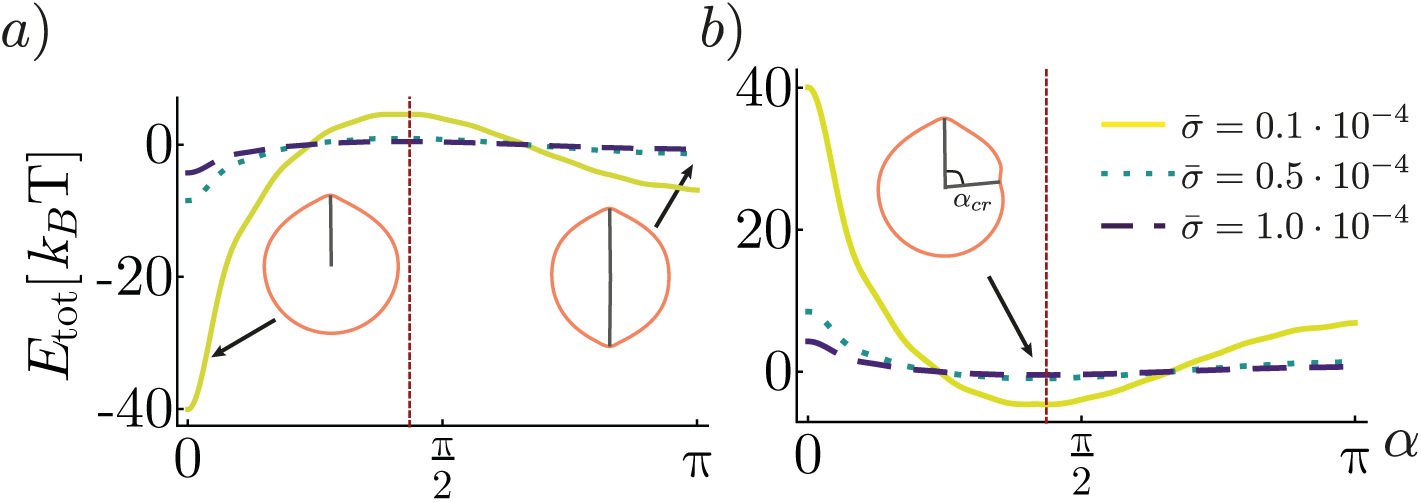
Total interaction energy for two point forces. The parameter *s* determines whether both forces are pushing (pulling) or one is pushing and the other pulling. **a)** *s* = 1. Both forces are pushing (pulling) and they experience strong short range attraction and weak long range repulsion. The critical angle determines weather the microtubules will coalesce or repel each other to the opposite poles. **b)** *s* = −1. In this case the forces have strong short range and weak long range repulsion, leading to a critical angle *α*_cr_ ≈ 0.45*π*. In both subplots the vertical red (dashed) line denotes *α*_cr_.

Combining eqns. 1 and 2 gives us an expression for the total energy of a vesicle, as a function of the magnitudes and positions of *n* point forces. The road from the general energy functional to the macroscopic energy function can be broken down into seven steps, which are detailed in the appendix.

Most importantly, we can derive a linearized shape equation, by using the Monge gauge (eqn. S13) and then minimizing the surface energy functional with respect to small deformations:

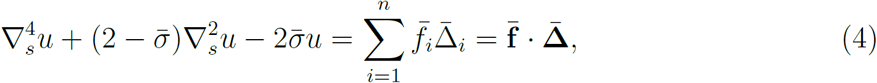

where *u*(*θ, ϕ*) is the relative deviation of the vesicle from a perfect sphere of radius *R* at position (*θ, ϕ*) as defined in eqn. S13. We also introduced the non-dimensionalized parameters 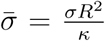 and 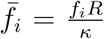. After finding the Green’s function of the left hand side (eqn. S25), its superposition gives the full solution of the inhomogeneous equation (eqn. S29), i.e. the deformation field *u* at any point (*θ, ϕ*). After a couple more steps we obtain the final energy function

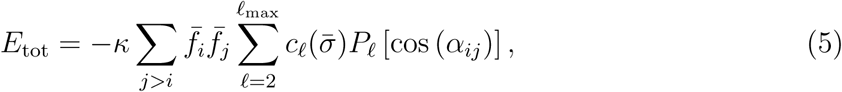

where *P*_*ℓ*_ denotes ℓ-th Legendre polynomial and cos(*α*_*ij*_) is the angle between the *i*-th and *j*-th microtubules at (*θ*_*i*_, *ϕ*_*i*_) and (*θ*_*j*_, *ϕ*_*j*_). The exact relation between these angles is given in eqn. S28. Finally 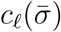 is an expansion coefficient determined by solving the shape equation. The upper cutoff *ℓ*_max_ corresponds to a high frequency (low wave length) cutoff which is justified by the fact that the membrane is not actually continuous but consists of lipids. In all of the proceeding analysis we used *ℓ*_max_ = 20.

## 3 Results and discussion

### 3.1 Analytic results for two microtubules

The system of two microtubules acting on a membrane can be treated fully analytically and can provide a valuable insight, for the *n >* 2 microtubule interactions. For simplicity we assume that both forces generated by the microtubules^1^ have the same magnitude 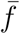. Then eqn. 5 simplifies to

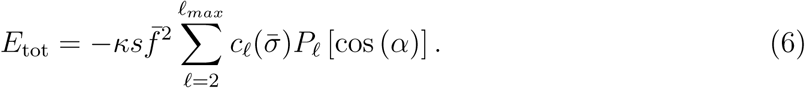

Where 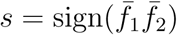 and *α* ∈ [0, *π*] is the angle between the two forces. Since *α* is the only configuration variable, we can easily plot the total surface interaction energy of the system. We do this because we are only interested in the interaction energy between the point forces and not in the actual energy of the vesicle surface. The exclusion of the “self-interaction energy”, which is the diagonal part in eqn. S31, does not mean that it is physically not measurable. It simply does not contribute to the rearrangement of point the forces. Note that the energy only depends on relative sign of the point forces *s* and not on each sign separately. Meaning that in our model it is the same when both point forces pull or push. This is a consequence of linearizion and would not hold true if the deformations *u*(*θ, ϕ*) become comparable with the vesicle size *R*.

In the case of two point forces of the same sign (both pushing or pulling) we can see a strong short range attraction together with a long range repulsion. This means that on a closed vesicle two point forces would either coalesce or repel each other until they reach the opposite poles of the vesicle, in both cases leading to emergent order (polar and nematic respectively). If the energy barrier at *α*_cr_ is too high (*E*(*α*_cr_) *− E*(*π*) ≫ *k*_*B*_*T*), the final state will depend on initial conditions. Forces that start close together will coalesce and forces that start further apart than *α*_*cr*_ will align at the opposite ends. In the case of one pulling and one pushing force we have only one ground state, in which the two point forces have a relative angle *α*_cr_ ≈ 1.4 rad.

Since the physical parameters of the system can be grouped into 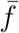 and 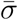, and *κ* is simply rescaling the energy, we can plot the full phase space of the system. Fig. 3 shows the magnitude of ⟨*α* ⟩, which is the expected equilibrium angle between two point forces in the system, for given pair of parameters 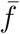 and 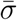. As we can see, for the most of the phase space ⟨*α* ⟩ = 0, meaning that the two point forces are coalesced in the equilibrium. However for small enough forces (or stiff enough membranes) 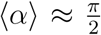. In this disordered state the value of *α* is essentially random and the average is 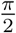 due to symmetry. Unfortunately the equilibrium average cannot predict the second (meta)stable state at *α* = *π*, since the latter is not a global minimum. The most interesting feature of figure 3, however, is that the shape of the plot is effectively independent of 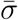. This intriguing property of the model is hiding in eqn. S26. If we expand 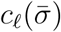 for large 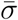 we get

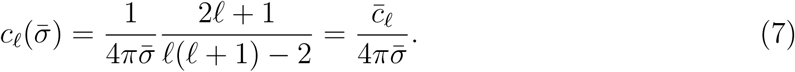

This expansion is justified since the biologically relevant cases have 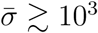. In fact for very small 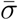 we see the shape of the phase space plot depends on both 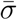 and 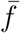. For large values of 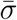 eqn. 5 simplifies to

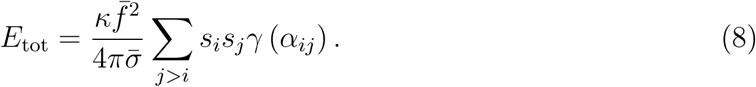

In 8 we again assumed for simplicity that all forces have the same magnitude, 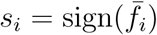 and we defined the two point interaction propagator

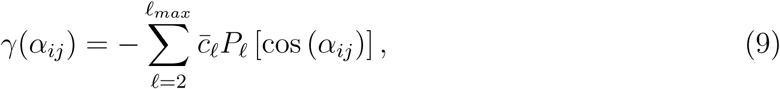

which contains the interaction part of the energy and is separated from the physical parameters of the model. Hence the nature of these interactions can be studied, without the need to consider specific values of the material parameters.

**Figure 3:**
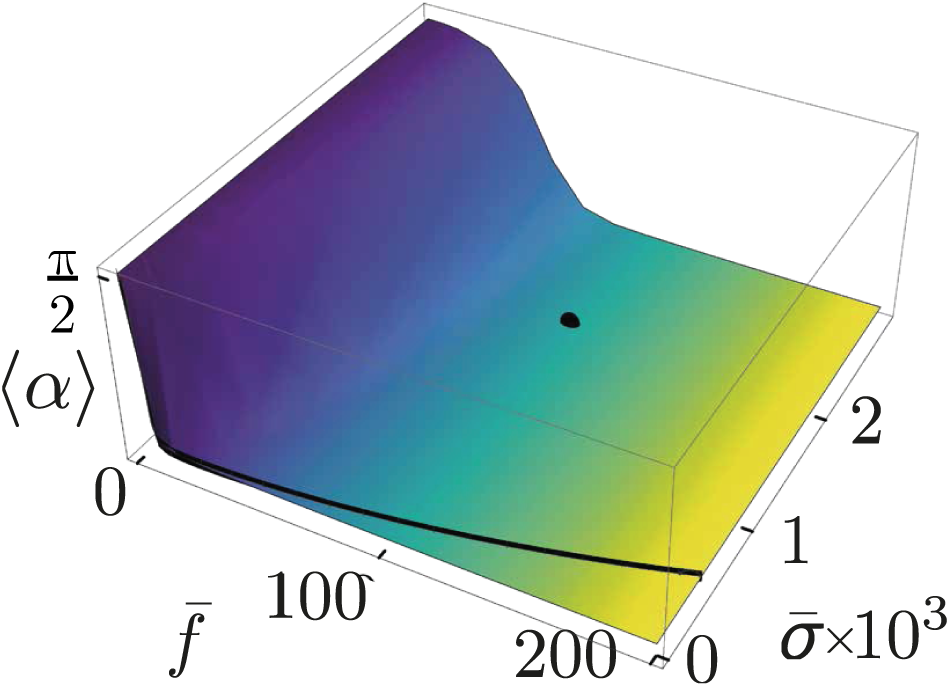
Expectation value of the equilibrium angle, between two point forces ⟨*α* ⟩, as a function of the magnitude of the force 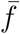 and the membrane tension 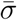. If the expected equilibrium angle between the point forces is 0 then they will coalesce. If 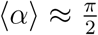 then the system is in a disordered state where the orientation of two point forces is random and the 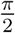 state is “winning” in the average by having the largest degeneracy. Black line denotes the boundary of the tether pulling region, 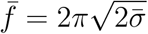. The black dot denotes the typical values used in our simulations, placing us firmly in the small deformation regime.

**Figure 4:**
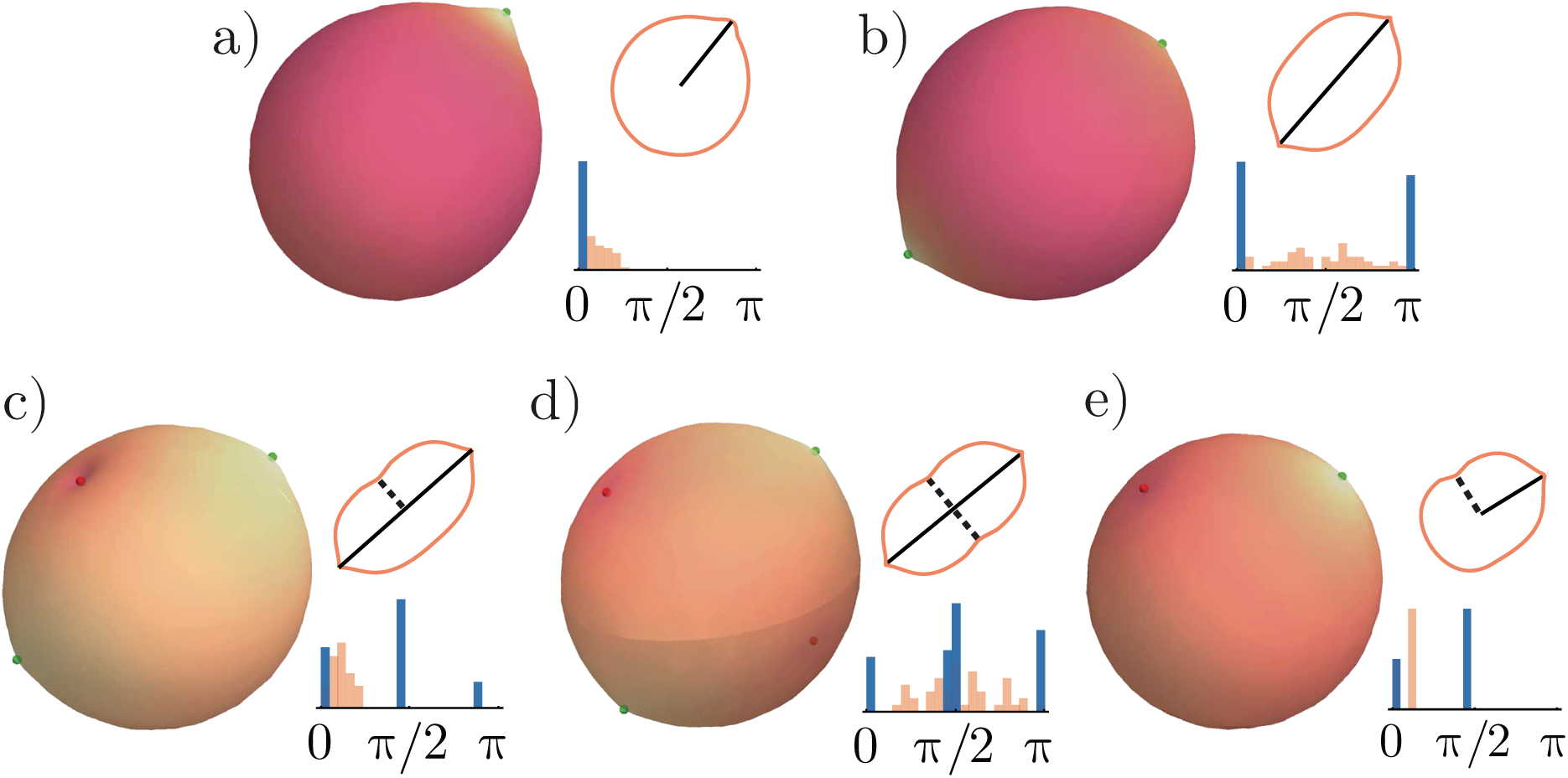
All five configurations of the final state of the Monte Carlo simulations of multiple point forces acting on the vesicle. Large 3D images are the actual plots of the final state of the simulation, accompanied by a sketch of the 2d cross section of the state (top) and a histogram (bottom) of the initial (light/orange bars) and final (dark/blue bars) distributions of the angles *α*_*ij*_ between the point forces. In all simulations we used the following parameter values: 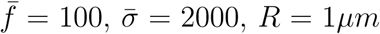 and *κ* = 10*k*_B_*T*. **a)** and **b)** 10 pushing microtubules. **c)** and **d)** 5 pushing and 5 pulling microtubules. **e)** 2 pushing and 1 pulling microtubules. **a)** Since all microtubules were close together initially they coalesced in one point. **b)** Here we have a broad initial distribution of microtubule positions consequently the final state has two points of bundled microtubules aligned at the opposite ends. **c)** Even if the initially all microtubules are close together, it is still likely that at least one of the flavors of microtubules (pushing or pulling) will create two poles of bundles. **d)** The most often observed configuration of the final state, where both flavors of microtubules are in nematic alignment. **e)** This state is very rare outside of 2 and 3 microtubule systems. Essentially for both flavors to be in polar alignment the initial conditions need to be perfectly tuned. Here we achieve this by having only one pulling force, hence depriving that flavor the possibility to make a nematic pattern and start with all microtubules close together such that the pushing ones are more likely to coalesce.

As we stated in section 2, our analysis relies on the small deformation assumption. In order to remain in this regime we use forces throughout our analysis which are significantly below the force 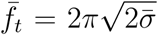 needed to stabilize a tether pulled from a vesicle, though the actual force needed to pull the tether is roughly 13% greater than this, for a point force as shown in (21). It is also worth noting that in reality the forces acting on the membrane are not point like but rather distributed over a patch of finite size. Both experiments and theory show that in the case of such forces the barrier to pull a tether is much higher compared to 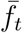 (22). This means that the regime of small deformations is not only a theoretically convenient approximation but rather a biologically relevant and stable geometric phase of a deformed vesicle. We use 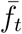 as our threshold for a “large” force (corresponding to a large deformation). In our analysis and simulations we use a micron sized membrane, i.e. *R* ≈ 10^3^ nm and material parameters and forces in typical range, *κ* ≈ 10 *k*_B_*T*, and *σ* ≈ 2 × 10^−2^ *k*_B_*T/*nm^2^ (23, Table 2). This leads to 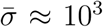. For an estimate of the force magnitude, we use the force that a typical cytoskeletal filament can generate (24, Ch.3), which is around 5 pN and translates into 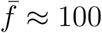. This is far below the tether force of 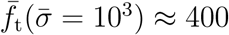.

### 3.2 Interactions between multiple microtubules

Eqn. 8 for the total energy of a system with two microtubules suggests that the *α*_*ij*_ dependent part of the equation is a generic function that does not depend on system parameters. To calculate the total energy of *n* point forces, we simply add the two point interaction propagator for all interacting pairs, thus we expect the extrema of *γ*(*α*_*ij*_) to still be present in the total interaction term, which we name Γ

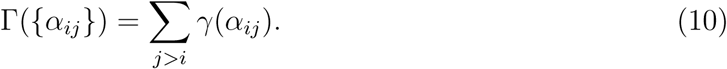

This means that the minimal energy states which are present in the *n* point force system are the same states as in the two point force system. Technically eqn. 10 does not prohibit the existence of additional energy minima. We therefore explore the state space of *n* = 8 − 16 microtubules using a Monte Carlo method. The results strongly suggest that there are no new stable configurations. The five configurations presented in fig.4 are the straightforward combinations of the three equilibrium configurations seen in fig.2.

To convince ourselves that the two point force analysis is also a good quantitative predictor of *n* point interactions we look at the dependence of the final configuration on the initial conditions. The prediction of eqn.8 is that each point force has a cone of attraction around it with an opening angle of *α*_*cr*_. If two point forces (of the same kind, i.e, both pushing or both pulling on the membrane) fall into each other’s cones of attraction they are likely to coalesce and if they fall outside of each other’s cones they are more likely to end up at the opposite poles. To this end we ran several Monte Carlo simulations where we placed all particles inside a cone of opening angle *α*_cone_ at the beginning and then let the system converge against a final state. The difference in numbers of aligned pairs versus anti-aligned pairs can be estimated by a quantity proportional to the average relative angle ⟨*α*_*ij*_⟩. We hence define a polar order parameter

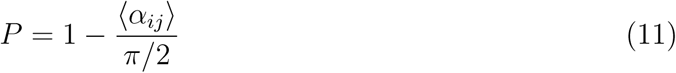

which is one in the polar phase i.e. when ⟨*α*_*ij*_⟩= 0, and it is zero in the nematic and disordered phases when 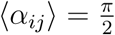. Hence to truly distinguish between the polar and nematic ordered states we need a second parameter *D* which measures the level of noise in the system and acts as a “disorder” parameter. As such (*P* = 1, *D* = 0) and (*P* = 0, *D* = 0) states correspond to polar and nematic orders respectively. The definition of *D* can be found in appendix B. In Fig. 5 we plot the dependence of the polar order parameter *P* on the opening angle of the initial state bounding cone and see that with growing cone angle *α*_cone_ we find a transition between polar and nematic end-states. This transition happens in the vicinity of *α*_cr_ as predicted from two point interaction analysis. Thus our point that *n >* 2 behavior does not qualitatively differ from two microtubule interactions is supported by this statistical analysis of the Monte Carlo simulations.

**Figure 5:**
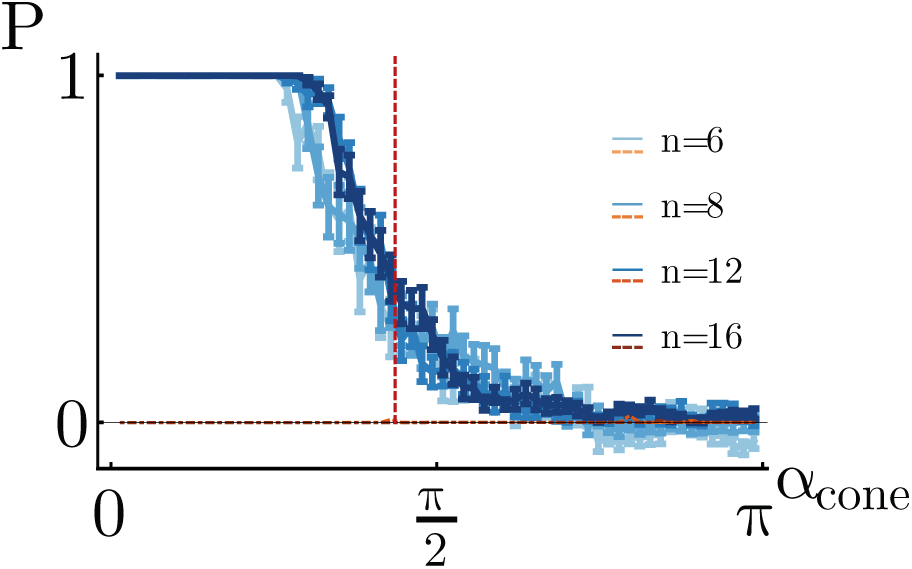
The dependence of the polar order *P* parameter on the opening angle of the cone of initial configurations *α*_cone_, which represents the maximum mutual angle, any two point forces can have at the beginning of the simulation. Each point is an average over several runs (the number varies between 10 and 30 Monte Carlo runs depending on *α*_cone_). The error bars display the sample standard error. Different shades of blue (solid with error bars) and orange (horizontal dot-dashed) lines depict the polar order and disorder parameters respectively, and the shade of the color indicates the amount of point forces pushing on the membrane. In all simulations we used the following parameter values: 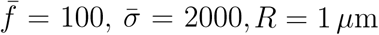 and *κ* = 10 *k*_B_*T*. The vertical red (dashed) line marks the position of the critical angle between two interacting point forces *α*_cr_ ≈ 1.4.

## 4 Conclusion

Together with actin and intermediate filaments, microtubules form an architecture that governs the shape of a cell, and therefore that of the plasma membrane surrounding it. The membrane, in turn, mediates the interaction between attached microtubules. Using analytical and numerical tools, we studied the effect of membrane mediated interactions on the rearrangement of microtubules. We found that force generating microtubules, when colliding with a deformable obstacle like a fluid membrane, can coordinate their growing state through the shape of distorted membrane between them. Our results suggest that the elastic properties of cellular membranes facilitate the bundling of microtubules. In particular, we showed that two vesicle-encapsulated microtubules attract each other for small angular separations and repel for large angles. As we demonstrated for up to 16 microtubules numerically and motivated analytically for any number, the outcome of collective interactions between multiple filaments is microtubule coalescence, which may be harnessed for protrusion formation (15, 25). Putting all the results together, our study suggests a possible mechanism underlying the preference of filaments for organizing in parallel configurations (26).

## Acknowledgments

AV was supported by the Netherlands Organisation for Scientific Research (NWO/ OCW), as part of the Frontiers of Nanoscience program and GD by the BaSyC Building a Synthetic Cell Gravitation grant (024.003.019) of the Netherlands Ministry of Education, Culture and Science (OCW) and the Netherlands Organisation for Scientific Research (NWO).

## A Derivation of macroscopic energy

In this part of the appendix we want to step through the derivation of the macroscopic energy. I.e. we want to show the steps that bring us from eqn.’s 1 and 2 to the eqn. 5. The total energy which is simply the sum of eqns. 1 and 2 reads as

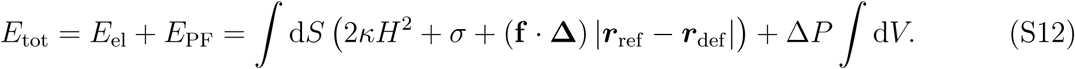

In the deformation term we used a coordinate free way to describe deviations from the reference state |***r***_ref_ − ***r***_def_|, where ***r***_ref_ and ***r***_def_ describe the membrane surface in the reference and deformed states respectively. To proceed we need to have a way to talk about the geometric quantities like *H* d*S* and V, this means that we need to choose a coordinate system and parametrize our membrane surface in that system, i.e. define a specific functional form of ***r***.

### A.1 Step 1: Monge gauge in spherical coordinates

We want to make use of the small deformation regime later to expand the surface energy and derive an analytically solvable shape equation. The natural parametrization for systems slightly deviating from a reference state is the Monge gauge. We use the spherical coordinates version of it to describe the membrane

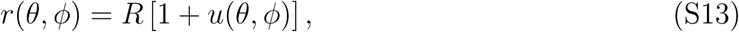

with *R* the radius of the undeformed (*reference*-)sphere and *u*(*θ, ϕ*) the relative deformation field, with respect to that reference-sphere. It is worth noting that this description itself is very general and can describe almost all shapes (apart from the ones with overhangs^2^). The small deformation regime is an additional assumption of

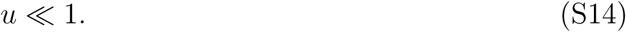

For this parametrization we can see how the magnitude of the deformation becomes:

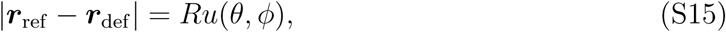

as used in the main text.

### A.2 Step 2: second order expansion in energy

Assuming small deformations, we expand *H*^2^ and 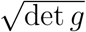 up to the second order, and *g*_*µν*_ = *∂*_*µ*_***r*** · *∂*_*ν*_***r*** and 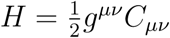, and

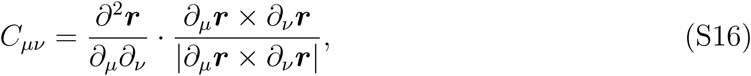

is the curvature tensor, also known as the 2nd fundamental form, with *µ, ν* ∈ {*θ, ϕ*}. In order to reliably perform this expansions we used Mathematica, which gave the following results^3^:

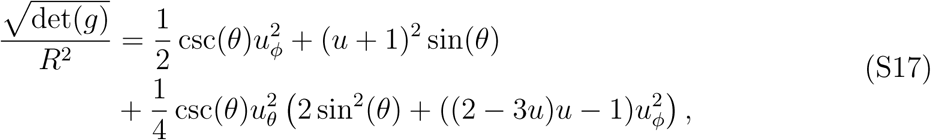

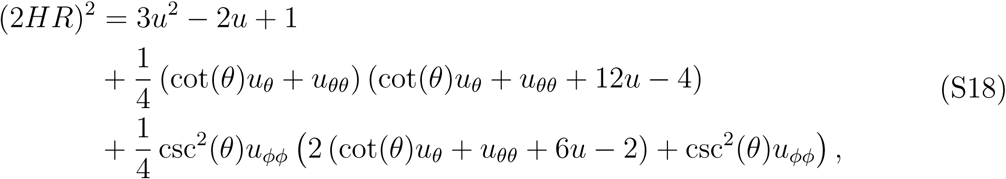

where we abbreviated the derivatives as indices i.e. *f*_*µ*_ = *∂*_*µ*_*f*. The expansions were performed up to the second order in *u* and all of its derivatives. Upon defining non-dimensionalized gradient and Laplacian operators in spherical coordinates, also known as surface gradient and Laplacian,

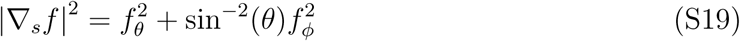

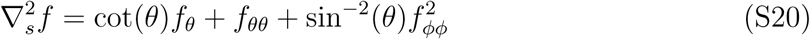

and reshuffling some terms we get

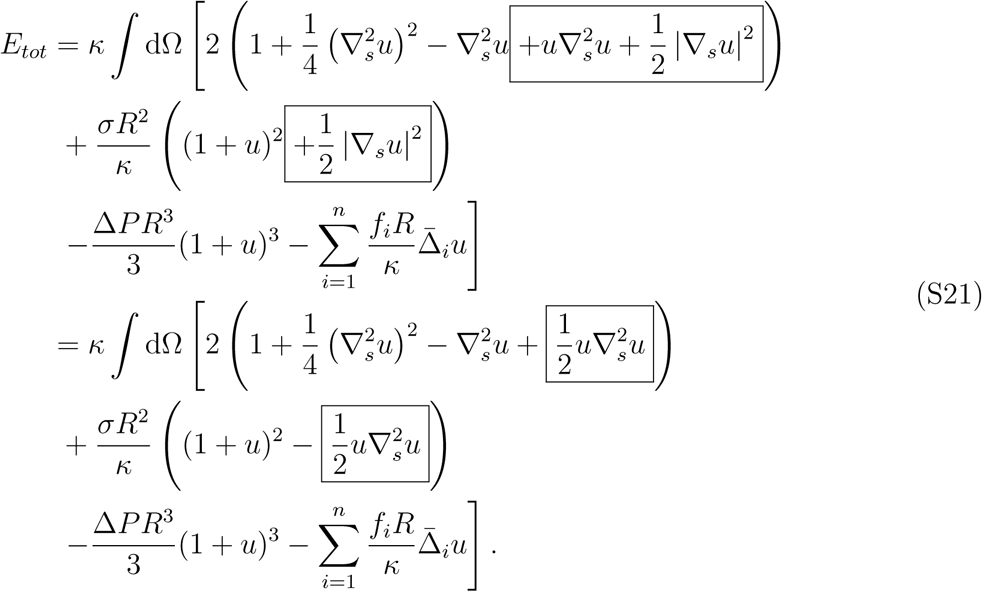

In the last equality sign we used partial integration to eliminate the absolute value signs

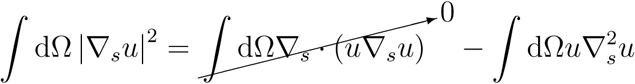

### A.3 Step 3: Energy variation

In previous step we arrived at a simplified energy functional in terms of a deformation field *u*(*θ, ϕ*). Next step is to find a surface configuration described by *u*(*θ, ϕ*) that minimizes this energy. For this end we vary the energy with respect to this field (i.e. *u* → *u* + *δu*), and further leverage the small deformation limit to replace Δ*P* with the Laplace pressure with the value for the sphere Δ*P* ≈ 2*σ/R* we get a shape equation.

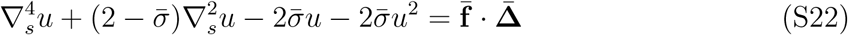

### A.4 Step 4: Linearization

Upon throwing away the nonlinear term in the shape equation S22, we arrive at eqn. 4.

### A.5 Step 5: Green’s function

To solve the eqn. 4 we first find its Green’s function. This is a function *G*(*θ* − *θ*_*i*_, *ϕ* − *ϕ*_*i*_) that solves the equation:

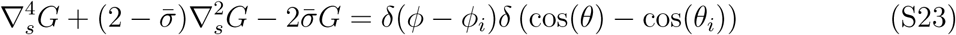

After expanding the delta functions in terms of spherical harmonics

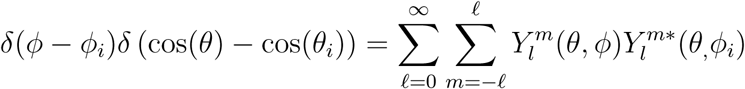

where ‘*’ denotes complex conjugate, we get a solution

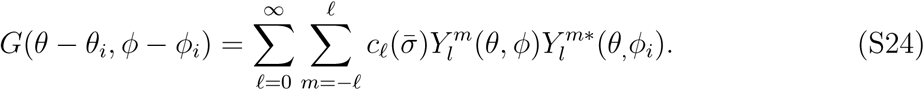

The first two modes in the expansion however need to be excluded. Since the zeroth mode (*ℓ* = 0) corresponds to the motion of the center of mass, which is irrelevant for our purposes and the first mode (*ℓ* = 1) corresponds to the conservation of the volume which is already accounted for in the energy functional with the Δ*P* term. Furthermore we use the addition theorem for spherical harmonics to simplify the second sum to *ℓ*-th degree Legendre polynomial, and finally arrive at

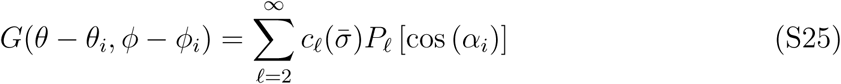

with the summand coefficients given by

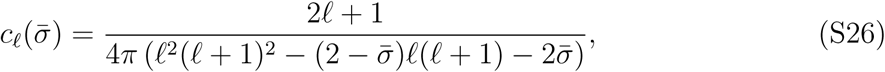

and *α*_*i*_ defined as the angle between the *i*-th deforming point force and and any other point on the membrane given by *θ* and *ϕ*.

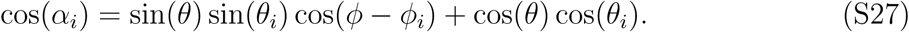

If this other point happens to be a position of *j*-th point force, then we get

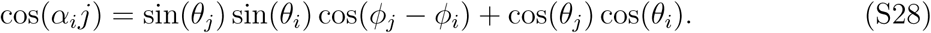

This makes *G*(*θ* − *θ*_*i*_, *ϕ* − *ϕ*_*i*_) a propagator of deformation from (*θ*_*i*_, *ϕ*_*i*_) to (*θ, ϕ*).

### A.6 Step 6: General solutions

The superposition of Greens’ functions gives the deformation field *u*:

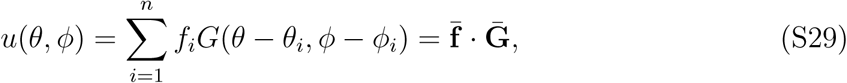

and hence

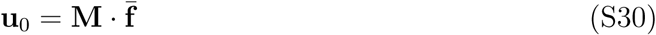

Where *u*_0*i*_ = *u*(*θ*_*i*_, *ϕ*_*i*_) is the vector of deformations at positions (*θ*_*i*_, *ϕ*_*i*_) and *M*_*ij*_ = *G*(*θ*_*i*_ − *θ*_*j*_, *ϕ*_*i*_ − *ϕ*_*j*_) is the deformation interaction matrix. The diagonal elements *M*_*ii*_ are the self interaction terms and do not depend on the positions of the deformations.

Substituting *u* back in the energy functional gives us the macroscopic surface energy which only depends on the position and magnitude of deformations

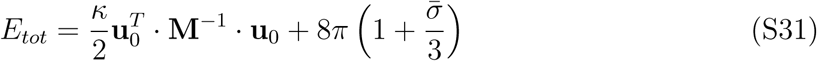

### A.7 Step 7: Legendre transformation

We use the Legendre transformation to transform **u**_0_ into its conjugate variable 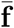, then **omit all constant**^4^ **terms** in the energy, use the symmetry of **M** and substitute the definition of the Green’s function from the eqn.S25

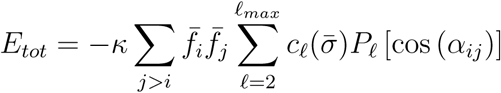

Actually *σ* can be either understood as penalizing the projected area of the vesicle, i.e. the energy cost per surface area to flatten out the thermal fluctuations of the vesicle. Or as a chemical potential giving the energy cost of incorporating new lipids in the membrane. The actual stretching of the membrane, i.e increase of the surface area per lipid is energetically so costly that the membrane will rapture long before that mode of deformation is accessed (23, Table 2). Point forces act along the radial direction of a spherical vesicle, i.e. 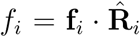. Here **f**_*i*_ is the *i*-th point force and 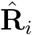 is the radial unit vector with spherical angles (*θ*_*i*_, *ϕ*_*i*_).

## B Disorder parameter

To analyze the results of the Monte Carlo simulations we introduced the polar order parameter *P* in eqn. 11. Unfortunately *P* = 0 is ambiguous since it could correspond to either nematic order or disorder. To distinguish between these states we defined a disorder parameter

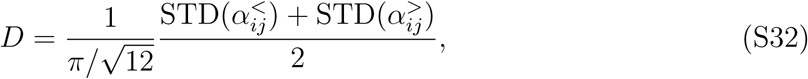

where the prefactor 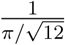 is the standard deviation of an uniform distribution on the interval [0, *π*] and serves as a normalization factor. STD stands for the standard deviation of a sample and 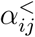 and 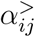 are the mutual angles in the final state that are smaller or greater than *π/*2. They can be formally defined as

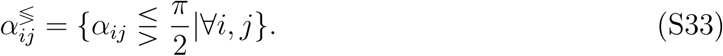

This parameter measures how narrow the peaks of the final distribution of *α*_*ij*_ are around the states 0 and *π*. Broad peaks would mean microtubules that did not fully coalesce, i.e., a disordered final state.

## Footnotes

1 From now on we will refer to the forces generated by the microtubules simply as point forces.

2 In 3D this would be a shape enclosing a volume which does not contain a point that can be connected to all points on the surface with a straight line that crosses this surface only at that point.

3 this equations are directly exported form MATHEMATICA and are hence not in the most readable form.

4 Constant with respect to the position of deformations {(*θ*_*i*_, *ϕ*_*i*_)}.

## References

[1] Fletcher, D. A., and R. D. Mullins. 2010. Cell mechanics and the cytoskeleton. Nature. 463:485.

[2] Revenu, C., R. Athman, S. Robine, and D. Louvard. 2004. The co-workers of actin filaments: from cell structures to signals. Nature reviews. Molecular cell biology. 5:635.

[3] Lodish, H., A. Berk, S. L. Zipursky, P. Matsudaira, D. Baltimore, and J. Darnell. 2000. The actin cytoskeleton. WH Freeman.

[4] Ridley, A. J. 2006. Rho gtpases and actin dynamics in membrane protrusions and vesicle trafficking. Trends in Cell Biology. 16:522–529.

[5] Le Clainche, C., and M.-F. Carlier. 2008. Regulation of actin assembly associated with protrusion and adhesion in cell migration. Physiological Reviews. 88:489–513.

[6] Fygenson, D. K., J. F. Marko, and A. Libchaber. 1997. Mechanics of microtubule-based membrane extension. Physical Review Letters. 79:4497.

[7] Emsellem, V., O. Cardoso, and P. Tabeling. 1998. Vesicle deformation by microtubules: a phase diagram. Physical Review E. 58:4807.

[8] Mogilner, A., and B. Rubinstein. 2005. The physics of filopodial protrusion. Biophysical Journal. 89:782–795.

[9] Atilgan, E., D. Wirtz, and S. X. Sun. 2006. Mechanics and dynamics of actin-driven thin membrane protrusions. Biophysical Journal. 90:65–76.

[10] e Silva, M. S., J. Alvarado, J. Nguyen, N. Georgoulia, B. M. Mulder, and G. H. Koenderink. 2011. Self-organized patterns of actin filaments in cell-sized confinement. Soft Matter. 7:10631–10641.

[11] Mesarec, L., W. Góźdź, S. Kralj, M. Fošnarič, S. Penič, V. Kralj-Iglič, and A. Iglič. 2017. On the role of external force of actin filaments in the formation of tubular protrusions of closed membrane shapes with anisotropic membrane components. European Biophysics Journal. 46:705–718.

[12] Kerssemakers, J. W., E. L. Munteanu, L. Laan, T. L. Noetzel, M. E. Janson, and M. Dogterom. 2006. Assembly dynamics of microtubules at molecular resolution. Nature. 442:709.

[13] Howard, J., and A. A. Hyman. 2003. Dynamics and mechanics of the microtubule plus end. Nature. 422:753.

[14] Liu, M., V. C. Nadar, F. Kozielski, M. Kozlowska, W. Yu, and P. W. Baas. 2010. Kinesin-12, a mitotic microtubule-associated motor protein, impacts axonal growth, navigation, and branching. Journal of Neuroscience. 30:14896–14906.

[15] Svitkina, T. M., E. A. Bulanova, O. Y. Chaga, D. M. Vignjevic, S. Kojima, J. M. Vasiliev, and G. G. Borisy. 2003. Mechanism of filopodia initiation by reorganization of a dendritic network. Journal of Cell Biology 160:409–421.

[16] Conde, C., and A. Cáceres. 2009. Microtubule assembly, organization and dynamics in axons and dendrites. Nature reviews. Neuroscience. 10:319.

[17] Dommersnes, P. G., and J.-B. Fournier. 2002. The many-body problem for anisotropic membrane inclusions and the self-assembly of saddle defects into an egg carton. Biophysical Journal. 83:2898–2905.

[18] Vahid, A., and T. Idema. 2016. Pointlike inclusion interactions in tubular membranes. Physical Review Letters. 117:138102.

[19] Canham, P. B. 1970. The minimum energy of bending as a possible explanation of the biconcave shape of the human red blood cell. Journal of Theoretical Biology. 26:61–81.

[20] Helfrich, W. 1973. Elastic properties of lipid bilayers: theory and possible experiments. Zeitschrift für Naturforschung C. 28:693–703.

[21] Derényi, I., F. Jülicher, and J. Prost. 2002. Formation and interaction of membrane tubes. Physical Review Letters. 88:238101.

[22] Koster, G., A. Cacciuto, I. Derényi, D. Frenkel, and M. Dogterom. 2005. Force barriers for membrane tube formation. Physical Review Letters. 94:068101.

[23] Phillips, R. 2018. Membranes by the Numbers. In: Bassereau P., Sens P. (eds), Physics of biological membranes, Springer Nature Switzerland, 73–105.

[24] Milo, R., and R. Phillips. 2015. Cell biology by the numbers. Garland Science.

[25] Weichsel, J., and P. L. Geissler. 2016. The more the tubular: Dynamic bundling of actin filaments for membrane tube formation. PLoS Computational Biology. 12:e1004982.

[26] Liu, A. P., D. L. Richmond, L. Maibaum, S. Pronk, P. L. Geissler, and D. A. Fletcher. 2008. Membrane-induced bundling of actin filaments. Nature Physics. 4:789.

